# A cryo-tomography-based volumetric model of the actin core of mouse vestibular hair cell stereocilia lacking plastin 1

**DOI:** 10.1101/825737

**Authors:** Junha Song, Roma Patterson, Jocelyn F. Krey, Samantha Hao, Linshanshan Wang, Brian Ng, Salim Sazzed, Julio Kovacs, Willy Wriggers, Jing He, Peter G. Barr-Gillespie, Manfred Auer

## Abstract

Electron cryo-tomography allows for high-resolution imaging of stereocilia in their native state. Because their actin filaments have a higher degree of order, we imaged stereocilia from mice lacking the actin crosslinker plastin 1 (PLS1). We found that while stereocilia actin filaments run 13 nm apart in parallel for long distances, there were gaps of significant size that were stochastically distributed throughout the actin core. Actin crosslinkers were distributed through the stereocilium, but did not occupy all possible binding sites. At stereocilia tips, protein density extended beyond actin filaments, especially on the side of the tip where a tip link should anchor. Along the shaft, repeating density was observed that corresponds to actin-to-membrane connectors. In the taper region, most actin filaments terminated near the plasma membrane. The remaining filaments twisted together to make a tighter bundle than was present in the shaft region; the spacing between them decreased from 13 nm to 9 nm. Our models illustrate detailed features of distinct structural domains that are present within the stereocilium.

## Introduction

Our senses of hearing and balance depend on the mechanosensitive hair bundles of the inner ear’s sensory cells, the hair cells. A bundle protrudes from a hair cell’s apical surface and consists of actin-filled stereocilia, which are arranged in a staircase (Roberts et al., 1988, Gillespie and Müller, 2009, Fettiplace and Kim, 2014). Each stereocilium consists of a distal tip, a shaft, and a proximal taper region, with the plasma membrane enclosing a highly-crosslinked actin filament core. About 400 actin filaments are found in each mouse utricle stereocilium (Krey et al., 2016); they are thought to run uninterrupted (Corwin and Warchol, 1991) from the tip along the shaft, parallel to the stereocilium longitudinal axis, before they end near the plasma membrane in the taper region (Tilney et al., 1986). The most central filaments condense into a rootlet structure, which penetrates deep into the actin meshwork of the cuticular plate and provides a pivot point that anchors the stereocilia (Tilney et al., 1980). In conventional transmission electron microscopy images, the core of the taper region and the initial insertion of the rootlet shows very high contrast when stained with osmium tetroxide, indicating either a very high protein density, an unusually high affinity for osmium, or both.

Actin filaments are heavily cross-linked by plastin 1 (PLS1; fimbrin) (Sobin and Flock, 1983, Tilney et al., 1989, Daudet and Lebart, 2002), espin (ESPN) (Zheng et al., 2000, Sekerkova et al., 2004, 2006), and fascin 2 (FSCN2) (Shin et al., 2010, Chou et al.,2011, Hwang et al., 2015). Early transmission electron microscopy studies suggested that stereocilia cores had a paracrystalline crosslinker pattern with full crosslinker occupancy (DeRosier et al., 1980, Jacobs and Hudspeth, 1990, Hackney et al., 1993). Mice lacking PLS1 display an actin filament core that is better ordered when compared to wild type mouse stereocilia (Krey et al., 2016), and strongly resembles the hexagonally packed actin core of chick cochlea {Tilney et al., 1983, #54878} and chick utricle {Shin et al., 2013, #26941}.

While hair cells with damaged hair bundles show some capacity for repair or replacement (Robertson et al., 1980, Liberman and Dodds, 1987, Gale et al., 2002), the stereocilia actin core is generally thought to be very robust. How the actin core is maintained is not entirely clear. Fluorescence studies of transfected mammalian hair cells in culture indicated that tagged actin or tagged ESPN incorporate into stereocilia tips (Rzadzinska et al., 2004). These findings suggested a continuous treadmilling mechanism, where actin is polymerized at the stereocilia tip and depolymerized near the taper region membrane, thus resulting in a net movement of all actin-filaments and their crosslinkers towards the proximal end of the stereocilia (Rzadzinska et al., 2004). This model was challenged by a study that included multi-isotope imaging mass spectrometry, differential temporal expression of labeled actin proteins in vivo, and fluorescence recovery after photobleaching; these experiments showed that while protein turned over rapidly at stereocilia tips, most protein remained stationary for weeks in the shaft, arguing against treadmilling (Zhang et al., 2012). Moreover, McDermott and colleagues used fluorescence imaging of genetically encoded in intact zebrafish larvae ear and showed dynamic turnover of both fascin 2b and actin β throughout the hair bundle (Hwang et al., 2015), further questioning the treadmilling model of actin turnover in stereocilia.

Quantitative mass spectrometry analysis in combination with electron tomography of high-pressure frozen, freeze-substituted, and resin-embedded tissue samples provided an estimated inventory of stereocilia proteins in chick utricle and showed that actin crosslinkers were not as abundant and regularly spaced as expected for a paracrystalline array (Shin et al., 2013). In the taper region, peripherally located actin filaments ended in close proximity to the plasma membrane whereas the central actin filaments of the actin core extended through the rootlet region into the cuticular plate. In resin-embedded samples, the rootlets appear as dark structures, which prevents discrimination of internal features.

The actin core is connected to its surrounding plasma membrane via RDX near the taper region (Kitajiri et al., 2004, Pataky et al., 2004, Zhao et al., 2012). In addition, unconventional myosins also serve as actin-to-membrane connectors throughout the stereocilium (Hasson et al., 1997). For example, MYO6 localizes towards the proximal end of stereocilia (Hasson et al., 1997), MYO7A along the entire shaft (Morgan et al., 2016), and MYO3A, MYO3B, and MYO15A at stereocilia tips (Belyantseva et al., 2003, Schneider et al., 2006, Merritt et al., 2012). The location of MYO1C is controversial; it is either concentrated at the upper tip link insertion side (Garcia et al., 1998, Steyger et al., 1998), where adaptation is thought to occur, or is found throughout the stereocilia membrane (Belyantseva et al., 2005).

Electron cryo-tomography (cryo-ET) has emerged as a powerful method for examining macromolecular structures in cells in general (Baker et al., 2017, Oikonomou and Jensen, 2017, Hutchings and Zanetti, 2018) and the cytoskeleton in particular (Jasnin et al., 2013, Turgay et al., 2017, McIntosh et al., 2018, Sun et al., 2018). We have previously reported cryo-ET studies at ∼3-4 nm resolution of frozen-hydrated individual stereocilia, which were isolated by blotting them away from the sensory epithelia onto a poly-lysine-coated grid (Metlagel et al., 2019).

Here we describe a simplified volumetric model of the actin core in the tip, shaft and taper regions of stereocilia harvested from utricular sensory epithelia of murine *Pls1*^*-/-*^ mice. We present here an ultrastructural 3D analysis of unstained, frozen-hydrated samples in an unstained near-native state. Although actin filaments adopt a highly ordered 3D organization in the shaft region, we also found significant longitudinal gaps along the actin filaments, indicating that actin filaments do not run uninterrupted from taper to tip. Examining the actin-actin crosslinker distribution, we found that only a fraction of possible crosslinking sites were occupied. In the tapered region, actin filaments adopt a complex 3D organization; the central core of the actin-filament bundle twists along the filament axis, which leads to a compacted, dense rootlet structure.

## Results

While we collected and reconstructed 26 tomograms of vestibular stereocilia from *Pls1*^*-/-*^ mutant mice, our in-depth analysis was focused on four tomograms, two for the tip and shaft region and two for the taper region. Our quantitative results presented here stem from one tomogram each of the shaft and taper region and the conclusions derived from these data were confirmed by the other two tomograms. All *Pls1*^*-/-*^ stereocilia reconstructed and examined displayed a higher degree of order for the actin core when compared to wild-type samples (Krey et al., 2016, Metlagel et al., 2019), with Fourier analysis yielding an estimated 4.3 nm resolution for the *Pls1*^*-/-*^ density maps examined. This limited resolution prevented us from docking atomic models into the density maps, but allowed us to build simplified volumetric ball-and-stick models into the density map, and to determine filament and cross-link numbers, dimensions, curvature, distances, and spacings.

### Manual model building followed by refinement

We obtained cryo-tomograms of structurally intact stereocilia that were blotted from utricles of *Pls1*^*-/-*^ mice. Examination of a single ∼1 nm cryo-tomogram slice central overview of the tip and shaft region (Fig. 1A), as well as a 10 nm cryo-tomogram slab close-up of the shaft region (Fig. 1B), revealed regularly spaced filamentous densities, which become more easily visible when tilting the 10 nm slab by ∼75 degrees out of plane (Fig. 1C). Tilting of the density maps along the stereocilia axis made it much easier to detect the actin-filament axis and place a ball-and-stick starting model onto the map. Rather than placing individual models one-by-one for each filament into the density maps, we simultaneously placed up to 17 parallel strands with an original center-to-center spacing of 12.5 nm (Fig. 1D). Once we obtained by visual inspection an acceptable global fit for an individual actin model layer to the ∼10 nm density slab (Fig. 1D), we locally adjusted each of the filament models manually to be at the center of the observed density maps (Fig. 1E). Length adjustments of the models in the tip region (Fig. 1F) resulted in a first model for each actin filament layer (Fig. 1G). After such longitudinal-orientation model building, we validated our model by replacing each cross-sectional slice (Fig. 1H) by a slice that represents an average density over 30 nm of slices (Fig. 1I-J) at the respective position, which greatly improved the signal-to-noise ratio. We refined the position of each filament model for each 30 nm-cross-averaged slice (Fig. 1K), resulting in a hexagonally close-packing 3D bundle (Figure 1L-M). We cross-checked our manual model tracing using a semi-automated actin tracing algorithm (Sazzed et al., 2018), which further improved the manual tracing. This stereocilium is almost certainly not from the tallest row, as its prolate tip with an asymmetric actin structure suggests that a tip link pulled on one side of the tip.

**Figure 1.**
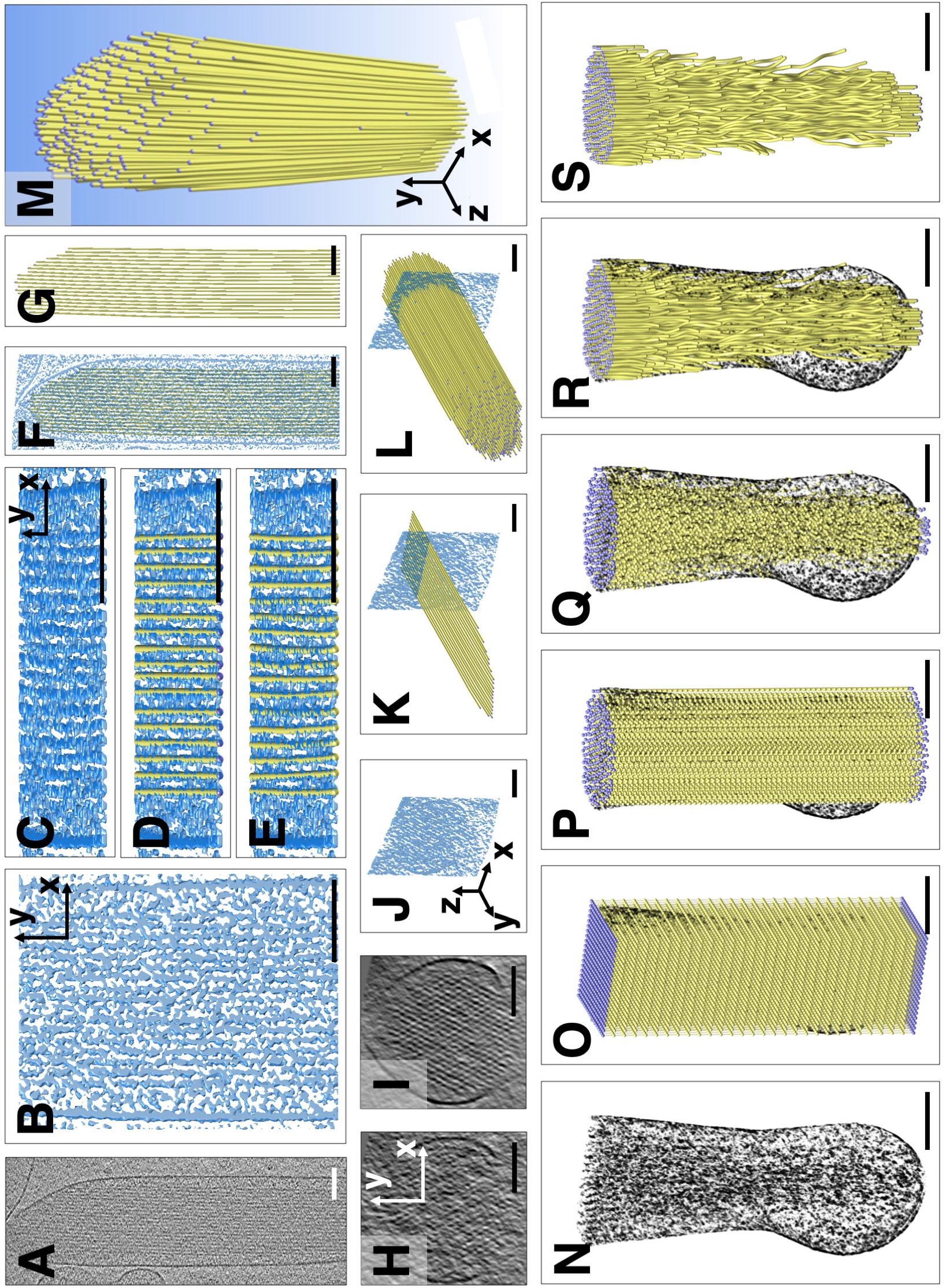
Volumetric model building of actin-filaments into tip and shaft, or shaft and taper region. (A) Cryo-tomographic grayscale map of a *Pls1*^*-/-*^ stereocilium; slice of ∼1 nm thickness in longitudinal (XY) orientation. (B) Solid rendering of a central portion of the shaft region, depicting a complex scenery of parallel actin filament and cross-linker densities. (C) Same shaft region as depicted in B, but rotated around the X-axis by 85 °, which allows to view almost precisely along the filament axis, making the filamentous nature of the densities more obvious. (D) Manual fitting of one layer of a simplified balls-and-sticks-model of parallel, tube-like actin filaments. (E) After global fitting for an individual model layer of parallel actin-filament tubes, the position of each ball was adjusted to achieve a local fit of each of the actin-filament tubes to their respective density. (F, G) Single layer model shown with (F) or without (G) a central slab density map. (H) Single slice of density map in cross-sectional (XZ) orientation. Note that the map is very noisy and periodicity is difficult to see. (I, J) A 30 nm Y-axis-averaged cross-sectional (XZ) slice showing clear periodicity of hexagonal packing of actin filaments in orthogonal view (I) and perspective (J) view. (K) Longitudinal single actin layer model superimposed onto single slice of a 30 nm-averaged cross-sectional map. Note that the model can be very accurately positioned into the density map shown in cross-sectional orientation. (L) The entire actin bundle model superimposed onto single slice of 30 nm-averaged cross-sectional map. (M) Entire model of actin filaments of the stereocilia actin core shown in perspective orientation. (N) Single ∼1 nm slice of cryo-tomographic 3D reconstructed volume in taper and rootlet region. (O) Rectangular prism containing 20 × 20 × 30 balls, with 20 × 20 balls spaced ∼12 nm apart in the cross-sectional plane of the taper region/rootlet and spaced 20 nm apart in the longitudinal stereocilia axis). (P, R) Corresponding balls along the filament axis were connected by sticks. Only balls inside a cylinder with a radius of the stereocilia membrane in the shaft region (top) were retained. (Q) Only balls corresponding to features of the density map inside the confines of the stereocilia membrane were retained. (S) Final coarse model of actin filament core, including the shaft, taper, and rootlet regions. Scale bars = 100 nm

For the tapered region of the stereocilia (Fig. 1N), we took a different approach, reflecting the fact that individual actin filaments in the bundle undergo a more complex path in the rootlet portion of density map. We started out by placing an array of 20 × 20 × 30 balls onto the corresponding density map (Fig. 1O). Corresponding balls were then connected along the filament axis. All model balls that fell outside the actin core density map were eliminated (Fig. 1P-Q). For each of the 30 cross-sectional layers spaced 20 nm apart we adjusted the position of balls using a 20-slice average at each of the positions (Fig. 1R).

Using our semi-automated filament tracing approach (Sazzed et al., 2018), we confirmed the manual model building approach and, in addition, carried out the fitting at smaller increments along the actin filament axis (Fig. 1S).

### Global bending of the actin filament core and actin filament gaps

Actin filaments in the stereocilia were parallel to one another throughout the shaft and tip region, and in the shaft region did not deviate from the stereocilia main axis (Fig. 2a). However, near the tip, all actin filaments deviated from the main axis towards the side of the lower stereocilium by 5 degrees on the side adjacent to a shorter neighboring stereocilium, and up to 8 degrees on the tall neighbor’s side (Fig. 2B). All filaments were bent in the same direction, which was obvious when viewing the volumetric ball-and-stick model head on (Fig. 2C). This curvature resulted in a displacement of the filament tips by ∼10-15 nm, which is hardly noticeable in longitudinal views (Fig. 2A), but becomes more obvious when viewed at an angle along the filament axis (Fig. 2B), or head-on (Fig. 2C). Note that all filaments underwent the same curvature and thus remained parallel to one another.

**Figure 2.**
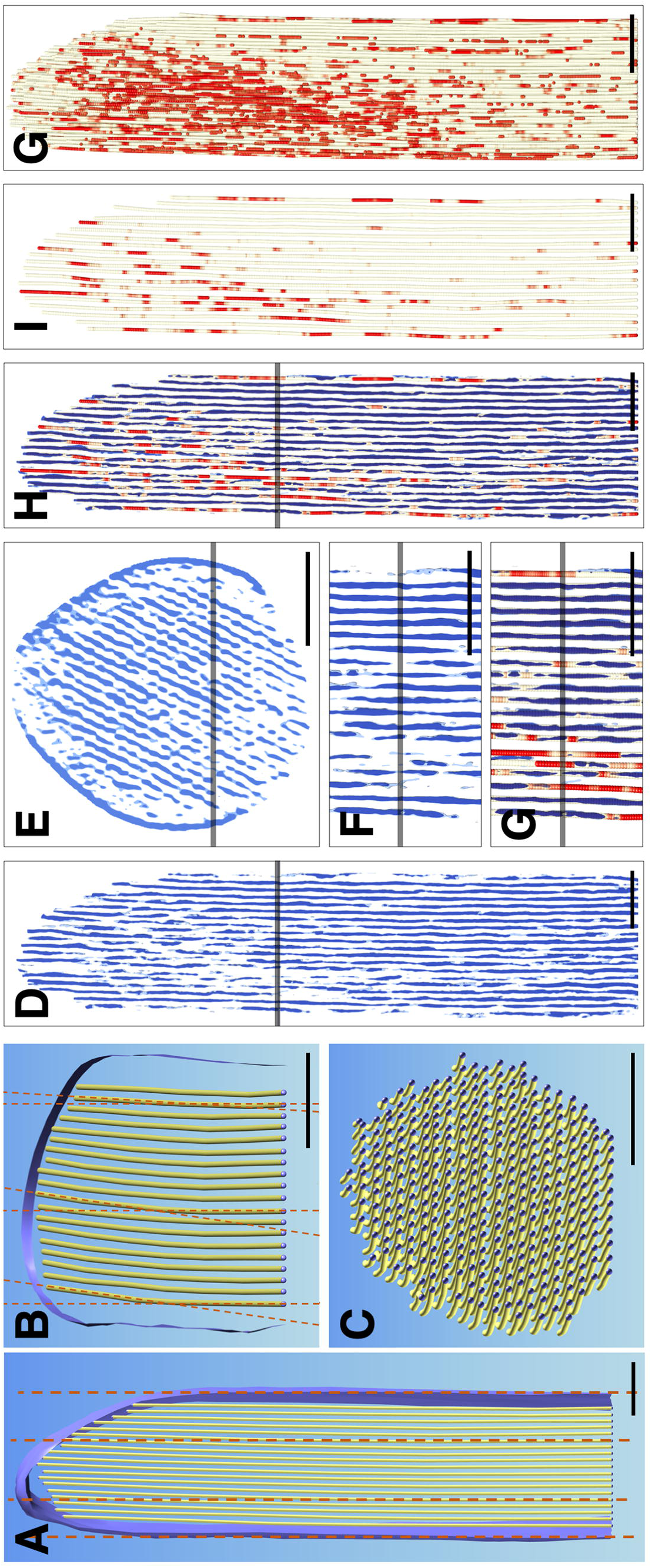
Actin core curvature and gaps. (A) Single central actin-layer model and plasma membrane surface rendering in longitudinal orientation. Dotted line indicates main axis of actin filaments and stereocilia. (B) Same single actin layer model as in A rotated around the X-axis to emphasize small deviations of 5-8 degree near the tip of stereocilia (C) En-face view of the actin filament model reveals that all actin filaments curve near the tip into the same direction, i.e., away from lower tip link insertion site. (D) Slab of 10 nm through 3D volume where each pixel inside the stereocilia membrane along the Y-axis has been replaced by a 30 nm Y-axis average to increase the signal-to-noise ratio in the direction of the filament. Note that significant gaps exist in the 30 nm averaged map. (E) Cross-sectional view of 30 nm averaged map rotated around X-axis. Note the periodicity of the density map and several gaps in the periodic pattern. (F) Close-up view of D. (G, H) Model of actin filament superimposed onto 30 nm averaged map. Care was taken to refine position of the model to reflect that 30 nm average map. Three map thresholds were chosen; we used cutoffs of ∼20% above, exactly at, and ∼20% below the average of the density map at the filaments and at the space between the actin filaments. Red color-coding reflects actual density value of the ball position with red being well below the average density, and thus a gap in the density map averaged for 30 nm along the actin filament axis. (I, J) Gap model (red) of actin filament core in single 2D layer (I) and the 3D bundle (J). Scale bars = 100 nm.

A thorough analysis of a map where each XZ slice was replaced by its 30 nm-slab average (Fig. 2D-H) revealed holes in the density map, both in longitudinal (Fig. 2D, 2F-H) and in cross-sectional (Fig. 2E) orientations. We overlaid our actin filament model onto the 30 nm averaged map and color-coded the model red in the gap regions where the density was missing (Fig. 2G). Gaps typically ranged in length from ∼20 to ∼75 nm, and are shown in Fig. 2I for an individual model layer or in Fig 2J for the entire actin filament core. There was no obvious pattern to the distribution of the gaps, and they were found throughout the stereocilium; there did appear to be more gaps near the tip of stereocilium, however (Fig. 2J).

### Membrane-to-actin connectors in tip and shaft regions

To better understand the interaction of actin filaments with macromolecules in the stereocilia tip region, we divided the density in our structure that lies between the top of the actin filaments (yellow lines) and the tip membrane (blue) into two regions, which we color-coded golden and maroon. Maroon-colored structures correspond to density within 10 nm of the distal end of the actin filaments, and thus constitutes the density map for proteins that may bind directly to actin filaments (Fig. 3A). In Fig. 3A, a 10-nm slab of the density that corresponds to a single actin model layer is shown. Fig. 3B shows the density as a 3D object in longitudinal orientation; the actin paracrystal was omitted in this view. When rotated 90° around the X-axis, one obtains an en-face view of the maroon-colored density that resides in close proximity to the actin filament ends. Note that there are a number of maroon-colored shapes of similar dimensions that are located near the actin filaments, whose ends are indicated by small dots (Fig. 3C, D). In Fig. 3D, the balls have a diameter of 6 nm, in accordance with the dimensions of actin filaments.

**Figure 3.**
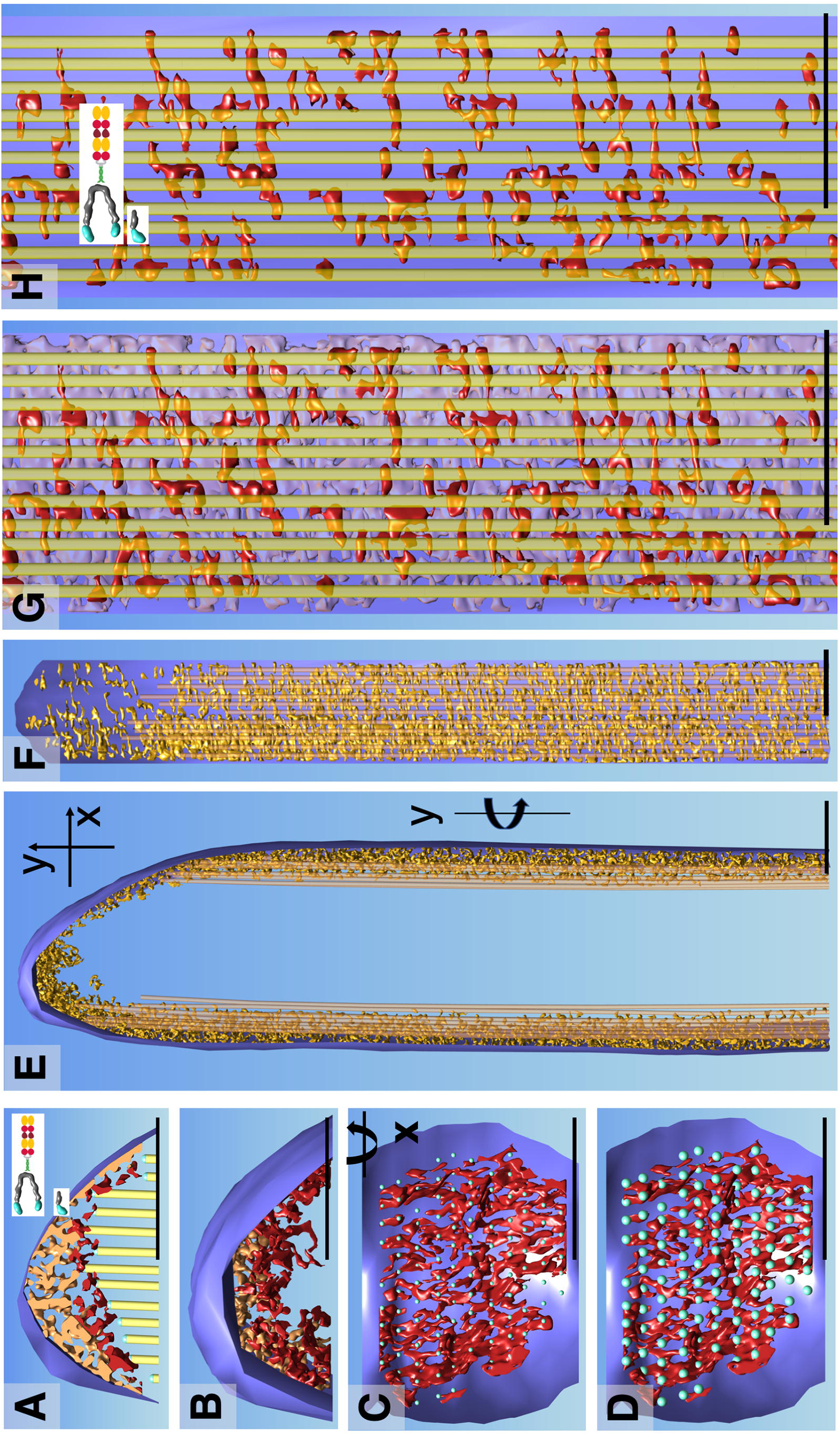
Actin-membrane crosslinkers in the tip and shaft regions. (A) Central density slab of the tip region with the most distal portion of the corresponding central actin-filament model layer. Note complex density near the lower tip link insertion site (left third of stereocilia tip), in stark contrast to the close proximity of the actin filament on the opposite stereocilia tip side. Red colored density rendering depicts map density within 10 nm proximity to the end of the actin filament. (B) 3D density corresponding to A. (C) En-face view onto the density map in 10 nm proximity to the actin filaments, showing a number of density lobes of similar size and what appears to be a non-random distribution. Small balls depict center location of each actin filament. (D) Same view as C but with blue balls representing the approximate actin filament diameter. Note that most but not all actin filaments show a corresponding red-colored density map. (E) Stereocilia in longitudinal orientation with the outermost layer of the actin filament of the corresponding circumferential membrane stretch of the entire stereocilia (∼one third). Density between outermost actin layer and membrane is depicted in golden color. (F) Corresponding en face view of the shaft region by rotating the right portion of E by 90 degrees around the Y-axis. (G, H) Close up views of F with red density corresponding to those in close proximity (10 nm) to the outermost actin filament layer and transparent purple density corresponding to density between the red density and the membrane plane. Note that the membrane is depicted as a single plane. (H) Outermost actin layer (yellow) and red density map within 10 nm proximity to the actin filaments. Note that the red densities are similar and size, shape and orientation with the densities seen in C and D and are consistent with models of unconventional myosins. Scale bars = 100 nm

In addition to connectors at stereocilia tips, we also studied the space between the actin filament core (yellow lines) and the plasma membrane (blue) in the shaft region (Fig. 3E-F). Our analysis was restricted to about one-third of the entire stereocilia membrane due to the missing wedge of data collection and the resulting data anisotropy. Fig. 3E shows the densities in the space between the plasma membrane and the most outer layer of actin filament in profile. Rotation around the Y-axis by 90° allows an en-face view of the density, which appears to be very complex. In order to simplify the scenery, we color-coded the density map within 10 nm proximity to the most outer actin filament layer (Fig. 3G-H). For a section of 115 nm x 400 nm, we counted ∼120 structures that are both close to the actin filaments and to the membrane (Fig. 3H), which extrapolates to ∼9000 actin-to-membrane connectors for a stereocilium of 5 µm. While the resolution is insufficient to determine molecular identity, many of the densities seen in Fig. 3H show similarity in size and shape to myosins visualized at similar resolution (Whittaker et al., 1995, Jontes and Milligan, 1997a, 1997b).

### Actin crosslinkers

To determine the 3D organization of the actin core, we created a simplified ball-and-stick model that represents actin filaments and crosslinkers. For quantitative analysis we focused on the layer direction in this hexagonally packed actin core, which was the least affected by the missing wedge-related data anisotropy. We examined the density map for three different stereocilia regions at a slab thickness of 10 nm, containing one layer of actin filaments (Fig. 4A-C). We adjusted the actin-filament model to best fit locally the density map. While we represented the actin filament as a round uniform cylinder, this is a gross simplification; the native actin filament is a helically-wound polymer, and at any point along the filament it has an elliptical cross-section, with periodically thicker and thinner densities.

**Figure 4.**
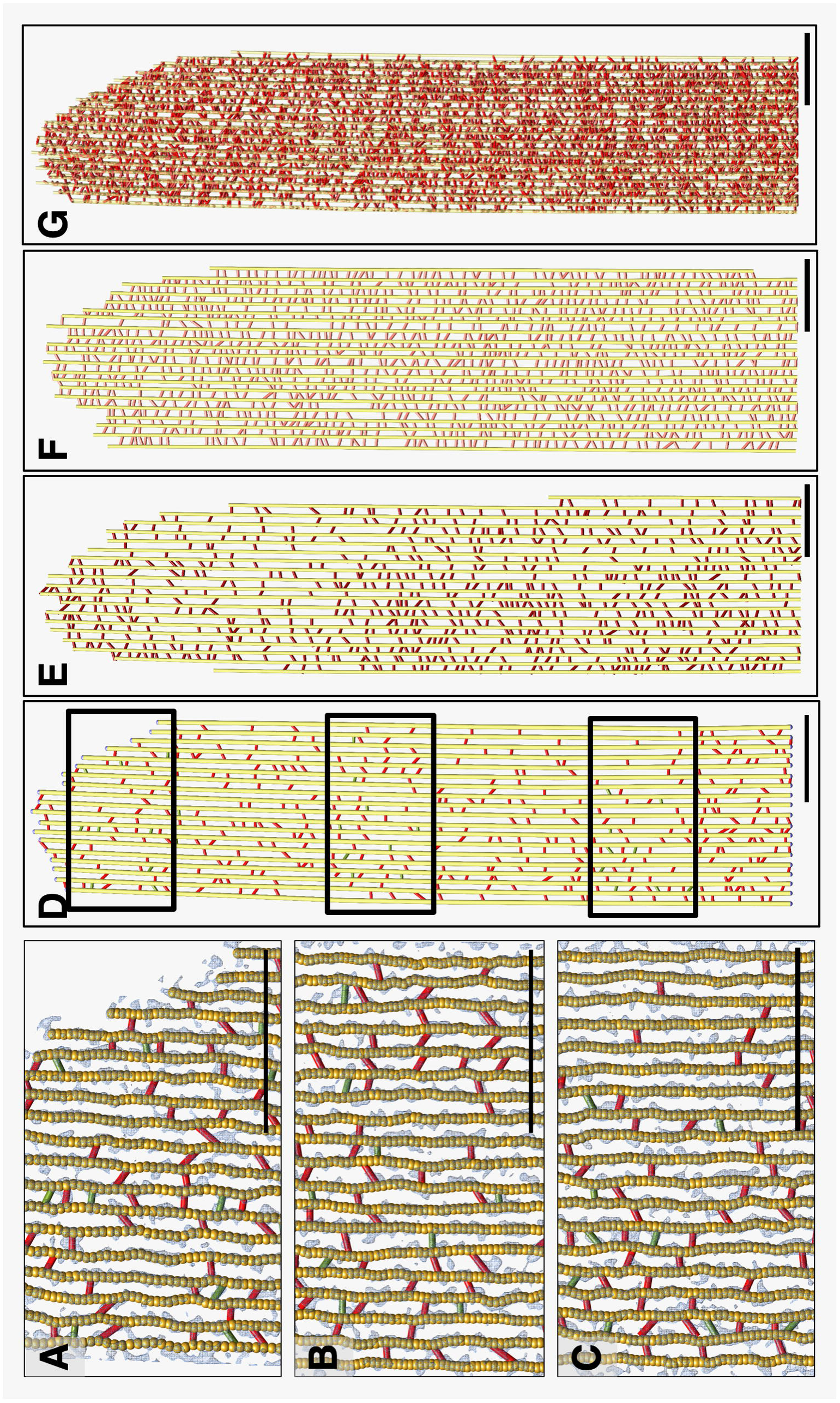
Actin-actin cross-linkers in all three longitudinal directions. Models showing a single actin filament plane in longitudinal orientation. (A-D) Actin-actin links in near-horizontal orientation. (E) Actin-actin links in a central plane oriented −60 degrees from the horizontal orientation. (F) Actin-actin links in a central plane oriented +60 degrees from the horizontal orientation. Note that this orientation is close to the Z-axis in data collection and thus along the path of the electron beam, where the densities are most affected by data anisotropy. Note the much higher density of crosslinkers as compared to the other two directions. (G) A ∼70 nm thick slab of the actin models and crosslinkers, which should resemble a projection TEM view though a ∼70 nm resin section. Note that there are some areas that appear more regular than others, whereas no obvious patterns are found in single actin filament model layers. Scale bars = 100 nm

Fig. 4D-F show three central planes of the resulting model. For the best oriented plane, which contained 19 actin filaments of 146 nm (four half turns) length, we found 47 actin-actin crosslinkers that were at multiples of 37 nm, as well as nine links that did not fit this pattern. Because of the elongation of the density map along the direction of the electron beam (data anisotropy), counting cross-linkers in the other two principle directions was less reliable and led to overestimates of the number of crosslinkers. Using the above crosslinker density and a total of 340 actin filaments (321 filament pairs), the extrapolated total number of crosslinkers is 78,000 for the 4.5 µm shaft of a prototypical stereocilium of 5 µm length. The 340 actin filaments, each of 4.5 µm, will comprise ∼554,000 actin monomers, yielding a crosslinker/actin monomer ratio of 0.14. The theoretical maximum is 0.23 crosslinkers/monomer (DeRosier and Tilney, 1982), which suggests a crosslinker occupancy of 61%. Fig. 4G shows a 70 nm thick slab of a model of actin core including the actin-actin crosslinkers.

Fig. 5A shows a 30 nm-averaged cross-sectional central slab through the stereocilium, which makes the hexagonal pattern of the actin core very obvious and allows the three main axes to be easily determined. In Fig. 5B, a corresponding model of a 10 nm central slab is shown with actin-filaments shown head on as yellow circles; red, firebrick red, and salmon colored crosslinkers signify actin-actin crosslinkers for each of the main three axes. A central portion of the three planes is shown in Fig. 5C. A single actin filament with all its crosslinkers to adjacent actin filaments is shown in various orientations in Figs. 5D-H. As is most obvious from Fig. 5G-H, there is no obvious pattern of clustering, and only a fraction of all possible positions for actin-actin crosslinking are occupied.

**Figure 5.**
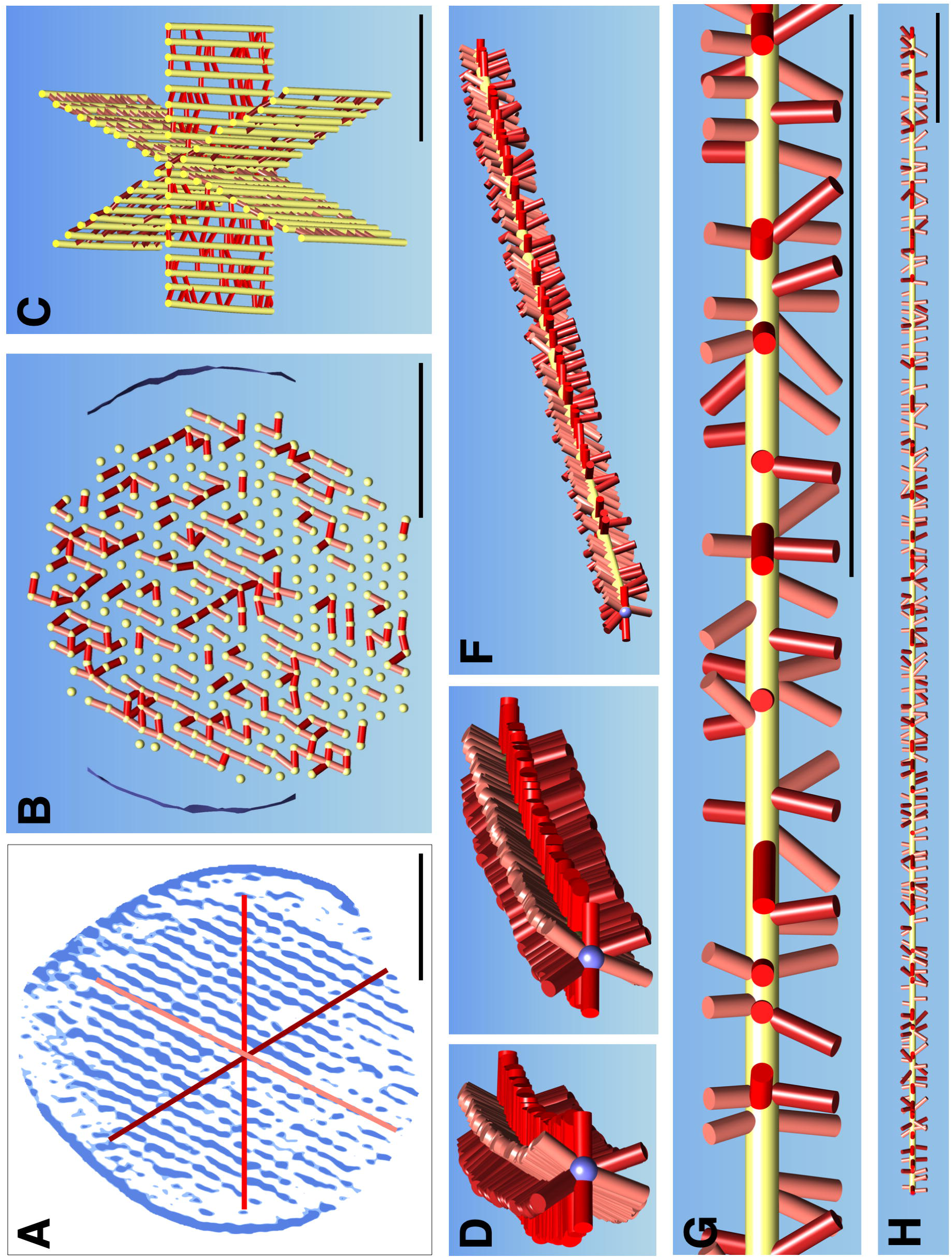
Actin-actin cross-linkers in cross-sectional directions and 3D model for one filament. (A) Cross-sectional view of 30 nm averaged map with the three main directions superimposed: horizontal plane, as well as plane at −60 degrees and +60 degrees are colored red, firebrick red and salmon, respectively. (B) Cross-sectional view of single actin filament layer (yellow dots) and corresponding actin-actin cross-connectors in XZ orientation. (C) Perspective representation of a small region of the actin cross-connectors in the three main regions. (D-H) Model representation of the actin filament component in the three main directions in different orientation (D-F) perspective views. (G) Close-up of a central region of the actin filament with its cross-connectors. (H) The actin filament (depicted in D-F) in its entire length with its associated cross-connectors. Scale bars = 100 nm.

### Actin 3D organization in the shaft regions

Filaments remained straight and therefore parallel to one another along the main stereocilia axis. We measured the actin-actin spacing at different locations of the actin core in the shaft region (Figs. 6A-D). The average spacing was 12.6 ± 1.2 nm (mean ± SD; n=2803), and did not differ at various positions along the stereocilium shaft (Fig. 6E). Distributions were fit well with a single Gaussian model, which indicates uniformity of crosslinking through the stereocilium.

**Figure 6.**
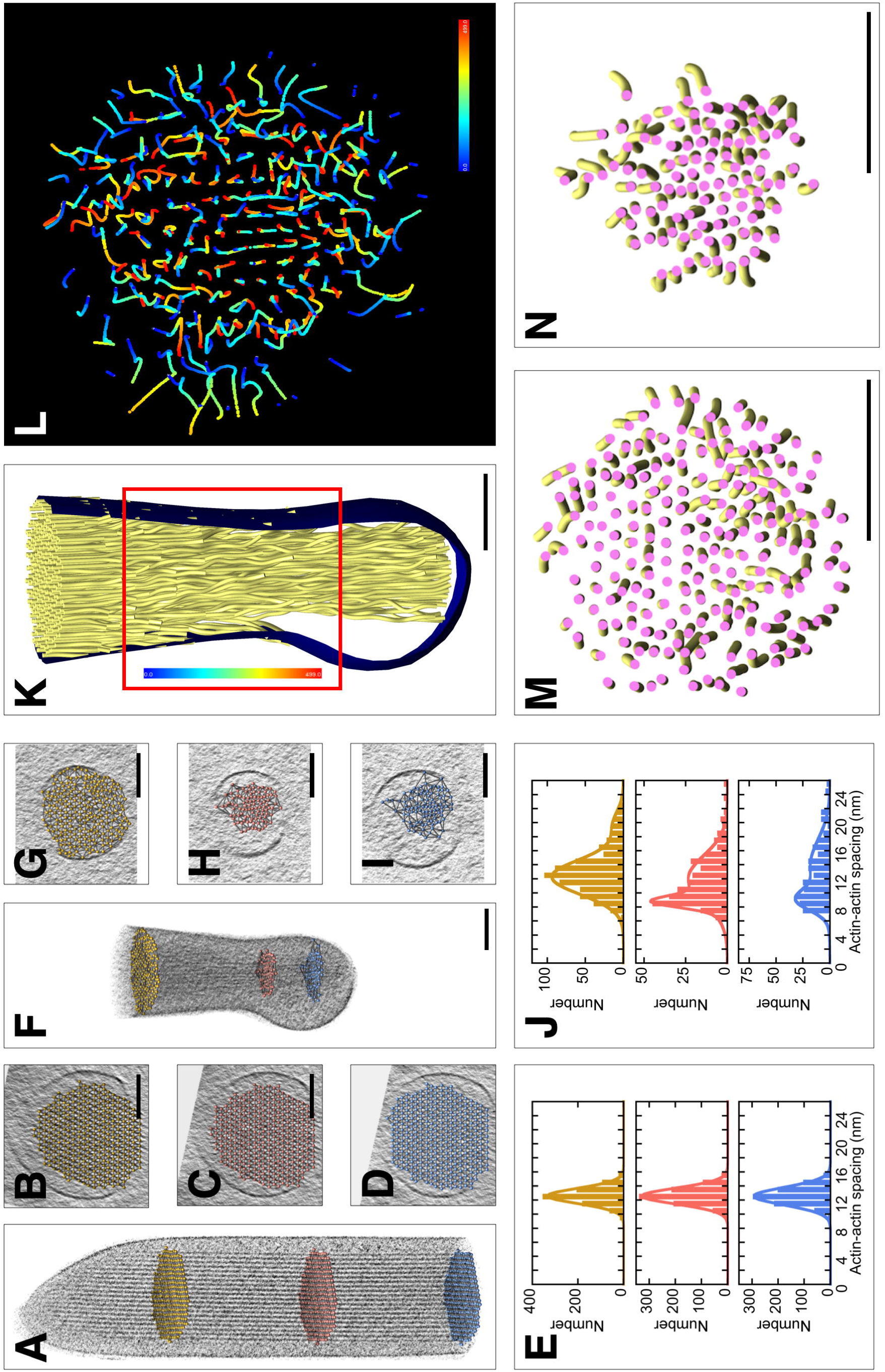
Changes in actin-actin spacing and 3D organization from shaft region to taper/rootlet region. (A-C) Stereocilia actin filament spacing in shaft region. (D-F) Stereocilia actin filament spacing in taper region. (A) Longitudinal orientation with balls in yellow, red, or blue at different locations of the shaft region. (B-D) Corresponding cross-sectional views with ball-model superimposed, allowing quantification of actin-actin spacing. (E) Histograms of actin-actin distance in shaft region. Single-Gaussian fits with peaks at 13.1 nm (yellow), 13.1 nm (red), and 13.1 nm (blue). (F) Longitudinal orientation with balls in yellow, red, or blue at different locations of the taper/rootlet region. (G-I) Corresponding cross-sectional views with ball model superimposed, allowing quantification of actin-actin spacing. (J) Histogram of actin-actin distance in taper/rootlet region. Note the change of spacing between shaft and rootlet region. Double-Gaussian fits with peaks at 12.8 and 20.0 nm (yellow); 9.2 and 12.5 nm (red); and 9.5 and 14.3 nm (blue). (K-L) Traces of actin filaments are they travel from the tapered region (blue) through the rootlet region (red). (M) Actin tube model in taper region. (N) Actin tube model in rootlet region. Note compaction of actin filaments in rootlet region and rotational twisting of actin filaments particularly around the core region. Scale bars = 100 nm.

### Actin 3D organization in the taper and rootlet regions

Fig. 1N suggests that our model contains not only the stereocilia taper, where the number of actin filaments is reduced before the stereocilium meets the hair cell soma, but also part of the rootlet, which normally extends into the soma and cuticular plate, anchoring the stereocilium to the cell (Furness et al., 2008). Examination of Fig. 1N shows that the membrane remains in close proximity to the actin core most of the distance to the stereocilium base, but abruptly disconnects from the cytoskeleton and forms a bubble around the last part. We suggest that the membrane is anchored to the stereocilium in the taper region (Tilney et al., 1986) but does not bind to the rootlet proper, which is fully intracellular and has no exposure to the plasma membrane (Furness et al., 2008).

Actin filaments in the taper region (Figs. 6F-I) adopted a more complex 3D organization than they did in the shaft. The spacing of actin filaments decreased from ∼13 nm near the shaft region to majority spacing of ∼9 nm in the rootlet (Fig. 6J). Many peripheral filaments end near the taper region membrane (Fig. 6K), whereas the inner actin core continues, with many but not all filaments adopting a curved or twisted trajectory (Figs. 6L-N), most apparent in a color-coded path-tracing of each actin filament (Fig. 6L). As can be seen in Fig. 6M-N, actin filaments transition from a loose organization in the taper region just above the rootlet (Fig. 6M) to a tightly packed arrangement in the rootlet region (Fig. 6N). The rootlet is osmiophilic, as is the central core of the stereocilium actin through the taper and up into the stereocilium shaft (Furness et al., 2008). Our density maps did not reveal any structural correlates of this central osmiophilic core beyond the compacting of actin filaments.

## Discussion

We focused here on the 3D structure of *Pls1*^*-/-*^ mutant mouse utricle stereocilia; a preliminary comparison of wild-type and knockout tomographic data sets revealed the higher order of the actin core in *Pls1*^*-/-*^ knockout stereocilia (Metlagel et al., 2019), which improved our model of the 3D organization of the actin core. The resolution in our tomograms was limited to 4.3 nm, however, which did not allow us to directly reveal the molecular identity of features visible in the density maps. Nevertheless, we were able to build simplified geometrical (volumetric) models, such as ball-and-stick models, into the density maps, which allowed us to determine the 3D organization and quantify the number of the actin filaments, actin-actin crosslinkers, and actin-membrane connectors.

Volumetric model building, first by manual fitting and then further refined by semi-automated fitting (Sazzed et al., 2018), resulted in actin models that ran parallel along the full length of the shaft and into the tip region. No abrupt changes in the actin filament orientation was observed. Accordingly, we averaged the map along 30 nm each of the actin filament models and thus replaced each voxel in the density map with a 30 nm average value. The resulting averaged map not only increased the signal-to-noise ratio and thus made the refinement of the actin model position much easier, but it also smoothed over fluctuations in map density caused by noise. A gap still visible in an averaged map therefore corresponded to a lack of density that extended over a significant length span, and therefore constitutes a gap in density. These gaps cannot be explained other than by breaks in the underlying actin filaments. Actin filaments throughout the shaft therefore run parallel to one another with a defined separation, yet display significant gaps along their lengths. These results suggest either that actin filaments are not formed continuously from taper to tip or that once the actin core has formed, it undergoes changes that include depolymerization of significant stretches. The gaps we encountered support the recent finding that actin turnover occurs throughout the stereocilium actin core (Hwang et al., 2015), and are not consistent with the treadmilling model for stereocilia actin turnover (Rzadzinska et al., 2004).

### Distinct structural features at stereocilia tips

We noticed several interesting features at stereocilia tips. For example, near tips, actin filaments deviated significantly from the main stereocilia axis in the direction opposite that of the tip link insertion site. While tip links were not present in the stereocilia model, the insertion site for the tip link can be readily recognized by the prolate shape of the tip, seen in all rows of stereocilia except for the tallest (Rzadzinska et al., 2004). This region of the stereocilium is subject to dynamic actin remodeling, even in adult animals (Zhang et al., 2012, Perrin et al., 2013, Narayanan et al., 2015), and actin’s structure at the tips is under the control of Ca^2+^ entering through transduction channels (Vélez-Ortega et al., 2017). While not conclusive, our results raise the possibility that transduction both stimulates local actin polymerization but also deflects filaments to avoid the site of local Ca^2+^ entry.

Density corresponding to proteins fills a gap between the end of the actin core and the plasma membrane; this distance is small opposite the putative tip link insertion, but can reach 40-50 nm where the tip link is presumed to insert. This protein density likely corresponds to the osmiophilic structure found underneath the tip link insertion, capping the actin core (Furness and Hackney, 1985). When under tension, the stereocilia membrane can be pulled away from this density by ∼15 nm (Assad et al., 1991), which was not seen in our model. Whether the density we observe includes tethers for the transduction channels (Powers et al., 2012), which could correspond to the gating spring (Corey and Hudspeth, 1983), is not known at present.

MYO3A, MYO3B, and MYO15A localize to stereocilia tips (Belyantseva et al., 2003, Schneider et al., 2006, Merritt et al., 2012). MYO15A in particular is thought to be deposited at high levels at the ends of the actin core (Belyantseva et al., 2003); in the shorter stereocilia rows, isoform 1 of MYO15A (MYO15A-L, the long form) is exclusively present and is found just underneath the tip link insertions (Fang et al., 2015). In the density maps corresponding to stereocilia tips, we observed structures that may correspond to unconventional myosins. Moreover, protein density at stereocilia tips could include the large N-terminal extension of MYO15A-L. Higher resolution is needed, however, to reveal their respective macromolecular identity.

### Actin-to-membrane connectors

We estimated that a 5 µm mouse utricle stereocilium has ∼9000 actin-to-membrane connectors, which could be RDX, myosins, or other proteins. While estimates for chick stereocilia were somewhat less, 5800-7300 per 5 µm (Shin et al., 2013), chick stereocilia are more narrow and hence have a smaller membrane circumference. Many of these connectors were tadpole-shaped, similar to how myosins appear at a similar resolution (Whittaker et al., 1995, Jontes and Milligan, 1997a, 1997b). Future high-resolution cryo-tomograms should provide the opportunity for docking high-resolution structures of the myosin motor domain on to the actin-to-membrane connectors seen along the stereocilia shafts.

Membrane-to-actin connectors are likely essential for maintenance of membrane tension in the stereocilia, which is important for controlling stereocilia shape (Prost et al., 2007) and transduction-channel gating (Powers et al., 2012, 2014, Peng et al., 2016). In most cells, the actin cytoskeleton and connectors that bridge it to the membrane are necessary to establish and control membrane tension (Pontes et al., 2017). Many membrane-to-actin connectors, including RDX and the myosin I family, bind strongly to PIP_2_, which is also essential for maintaining adhesion of the membrane to the cytoskeleton (Raucher et al., 2000). Notably, PIP_2_ is prominent in stereocilia (Hirono et al., 2004, Effertz et al., 2017). Actin-to-membrane connectors occupy far fewer than the theoretical maximum number sites on the periphery of the actin core, which may be essential for minimal impediments to transport of proteins along the shaft.

### Actin-actin crosslinkers

We also studied extensively the number and distribution of actin-actin crosslinker and they were similar to estimates previously made from our correlative study incorporating quantitative mass spectrometry and electron tomography of chick utricle stereocilia (Shin et al., 2013). In a model 5 µm long mouse utricle *Pls1*^*-/-*^ stereocilium, we estimated the presence of ∼78,000 crosslinkers for 554,000 stereocilia actin monomers in 340 actin filaments. These values compare to 60,000-90,000 crosslinkers and 400,000 stereocilia actin monomers in a chick stereocilium of the same length, albeit with 210 actin filaments (Shin et al., 2013). Targeted proteomics of CD-1 mouse stereocilia suggested the presence of 30,500 PLS1, 16,100 FSCN2, and 14,800 ESPN in a stereocilium of 400,000 actin monomers (Krey et al., 2016). Removing the PLS1 molecules (because we are modeling *Pls1*^*-/-*^ stereocilia) and extrapolating to the larger number of actin monomers in our model stereocilium, those mass spectrometry experiments predict a total of 45,000 non-PLS1 crosslinkers per stereocilium, reasonably close to the 77,000 we counted. Several actin crosslinkers increased in abundance, albeit not significantly, in targeted proteomics measurements comparing wild-type and *Pls1*^*-/-*^ stereocilia (Krey et al., 2016).

The uniformity of actin-actin spacing throughout the stereocilium suggests either that one crosslinker controls that spacing or that multiple crosslinkers have similar properties. Rigidity of actin-fascin crosslinks and flexibility of actin-espin crosslinks suggests that the uniform actin-actin spacing is set by ESPN crosslinkers rather than FSCN2 {Shin et al., 2009, #95951}, although they should be present at similar abundance.

We noticed that individual actin layers showed little sign of clustering of actin-actin crosslinkers. Together with the results from gap analysis, this results argues that the actin core’s 3D organization is more gel-like than paracrystalline, consistent with the dynamic nature of actin exchange within the core (Hwang et al., 2015).

As we noted in our preliminary study (Metlagel et al., 2019), the actin-actin spacing measured here (∼13 nm) is considerably larger than the ∼8 nm we measured previously for *Pls1*^*-/-*^ mutants (Krey et al., 2016). One of the major advantages of the electron cryo-tomography approach we used here is that we prepare samples by rapid freezing, with no fixation, dehydration, and staining steps that could distort stereocilia dimensions. For this reason, we believe that the 12.6 nm actin-actin spacing is an accurate estimate for actin cores crosslinked by a combination of FSCN2 and ESPN. In addition, this comparison also demonstrates how much tissue distortion conventional electron microscopy techniques introduce.

### Actin filaments in the taper region

We found that actin filaments in the taper region adopt a twisting path that results in the compaction of the filaments in the rootlet region. The compression we observed of the actin core corresponds to the absence of actin-actin crosslinkers previously reported for mouse utricle (Krey et al., 2016). Several membrane or membrane-associated proteins, including PTPRQ and RDX (Goodyear et al., 2003, Pataky et al., 2004, Zhao et al., 2012), are associated with the taper region, as is the protein TPRN (taperin), which could be binding protein for actin pointed ends (Rehman et al., 2010). It is not clear, however, what brings the actin filaments together. TRIOBP is a good candidate; it has been proposed to wrap around actin filaments in the rootlets (Kitajiri et al., 2010). Although our density maps do not provide evidence for a filamentous protein playing this role, higher-resolution data could provide evidence for TRIOBP and other actin-associated proteins. We believe that improvements in resolution will be accomplished by motion-correction of tomographic data sets and subsequent sub-tomogram averaging of repeated volume.

### Implications

We took advantage of the *Pls1*^*-/-*^ mutant mouse line, which allowed a more direct comparison to our previous work on chick utricle stereocilia (Shin et al., 2013). Moreover, the high order of the *Pls1*^*-/-*^ cytoskeleton (Krey et al., 2016) improved our ability to accurately develop our stereocilium model. This model improved substantially on our previous model for chick stereocilia, which used resin-embedding processing for transmission electron microscopy (Shin et al., 2013). In chick utricle stereocilia, which have low levels of PLS1 as compared to FSCN2, the dominant actin-actin crosslinker distance was ∼8 nm. This distance corresponds well to the predominant actin-actin spacing in mouse utricle stereocilia prepared from *Pls1*^*-/-*^ mutants using conventional processing, but is much smaller than the ∼13 nm measured here. Electron cryo-tomography using rapidly frozen samples, as we used here, provides the ability to see the cytoskeleton in near-native dimensions with little or no distortion. Future modeling of stereocilia structure will include those of wild-type mouse utricle, as well as stereocilia from inner and outer hair cells of the mouse cochlea.

## Materials and Methods

Blotting of stereocilia onto microscope grids, vitrification, cryo-tomographic data collection and 3D reconstruction have all been described in detail previously (Metlagel et al., 2019). In short, the sensory epithelium was blotted onto the lacey carbon support film of an EM grid, transferring intact stereocilia to the grid. Samples were vitrified using ultra-rapid plunge-freezing. Single axis cryo-tomograms were collected on a Krios TEM (Thermo Fisher) operated at 300 kV with a nominal defocus of 3.5-4.5 µm using a Falcon 2 camera in integration mode at 0.47 to 0.59 nm pixel size. Typical dose for single axis data collection was 80-100 electrons/Å^2^). Tomogram 3D volumes were reconstructed using IMOD (Kremer et al., 1996), using either weighted back-projection or the SIRT method (Agulleiro and Fernandez, 2011). To improve contrast, we filtered tomograms with recursive median or bilateral filtering in Priism (Chen et al., 1992) or IMOD. We used UCSF–Chimera software (Pettersen et al., 2004) for tomographic 3D volume visualization, model building and quantitatively analysis. Where mentioned in the results section we created a map where each stereocilia cross-section density map slice was replaced with a 30 nm average along that axis, which simplified manual and semiautomated tracing and trace refinement (Sazzed et al., 2018). Tracking of actin filaments in the taper/rootlet region was carried out with the assistance of Trackmate (Tinevez et al., 2017).

## Notes

#### Summary of Updates

Affiliations for Salim Sazzed and Jing He are corrected to Department of Computer Science, Old Dominion University

## References

Agulleiro, J. I. and Fernandez, J. J., 2011. Fast tomographic reconstruction on multicore computers. Bioinformatics. 27, 582–583.

Assad, J. A., Shepherd, G. M. G. and Corey, D. P., 1991. Tip-link integrity and mechanical transduction in vertebrate hair cells. Neuron. 7, 985–994.

Baker, L. A., Grange, M. and Grünewald, K., 2017. Electron cryo-tomography captures macromolecular complexes in native environments. Curr Opin Struct Biol. 46, 149–156.

Belyantseva, I. A., Boger, E. T. and Friedman, T. B., 2003. Myosin XVa localizes to the tips of inner ear sensory cell stereocilia and is essential for staircase formation of the hair bundle. Proc. Natl. Acad. Sci. USA. 100, 13958–13963.

Belyantseva, I. A., Boger, E. T., Naz, S., Frolenkov, G. I., Sellers, J. R., Ahmed, Z. M., Griffith, A. J. and Friedman, T. B., 2005. Myosin-XVa is required for tip localization of whirlin and differential elongation of hair-cell stereocilia. Nat Cell Biol. 7, 148–156.

Chen, H., Clyborne, W. K., Sedat, J. W. and Agard, D. A., 1992. Priism: an integrated system for display and analysis of 3-D microscope images. Proceedings of SPIE. 1660, 784–790.

Chou, S. W., Hwang, P., Gomez, G., Fernando, C. A., West, M. C., Pollock, L. M., Lin-Jones, J., Burnside, B. and McDermott, B. M., 2011. Fascin 2b is a component of stereocilia that lengthens actin-based protrusions. PLoS One. 6, e14807.

Corey, D. P. and Hudspeth, A. J., 1983. Kinetics of the receptor current in bullfrog saccular hair cells. J. Neurosci. 3, 962–976.

Corwin, J. T. and Warchol, M. E., 1991. Auditory hair cells: structure, function, development, and regeneration. Annu Rev Neurosci. 14, 301–333.

Daudet, N. and Lebart, M. C., 2002. Transient expression of the t-isoform of plastins/fimbrin in the stereocilia of developing auditory hair cells. Cell Motil Cytoskeleton. 53, 326–336.

DeRosier, D. J. and Tilney, L. G., 1982. How actin filaments pack into bundles. Cold Spring Harb Symp Quant Biol. 46 Pt 2, 525–540.

DeRosier, D. J., Tilney, L. G. and Egelman, E., 1980. Actin in the inner ear: the remarkable structure of the stereocilium. Nature. 287, 291–296.

Effertz, T., Becker, L., Peng, A. W. and Ricci, A. J., 2017. Phosphoinositol-4,5-bisphosphate regulates auditory hair cell mechanotransduction channel pore properties and fast adaptation. J Neurosci. pii: 1351–17. doi: 10.1523/JNEUROSCI.1351.

Fang, Q., Indzhykulian, A. A., Mustapha, M., Riordan, G. P., Dolan, D. F., Friedman, T. B., Belyantseva, I. A., Frolenkov, G. I., Camper, S. A. and Bird, J. E., 2015. The 133-kDa N-terminal domain enables myosin 15 to maintain mechanotransducing stereocilia and is essential for hearing. eLife. 4,

Fettiplace, R. and Kim, K. X., 2014. The physiology of mechanoelectrical transduction channels in hearing. Physiol Rev. 94, 951–986.

Furness, D. N. and Hackney, C. M., 1985. Cross-links between stereocilia in the guinea pig cochlea. Hearing Res. 18, 177–188.

Furness, D. N., Mahendrasingam, S., Ohashi, M., Fettiplace, R. and Hackney, C. M., 2008. The dimensions and composition of stereociliary rootlets in mammalian cochlear hair cells: comparison between high- and low-frequency cells and evidence for a connection to the lateral membrane. J Neurosci. 28, 6342–6353.

Gale, J. E., Meyers, J. R., Periasamy, A. and Corwin, J. T., 2002. Survival of bundleless hair cells and subsequent bundle replacement in the bullfrog’s saccule. J Neurobiol. 50, 81–92.

Garcia, J. A., Yee, A. G., Gillespie, P. G. and Corey, D. P., 1998. Localization of myosin-Iβ near both ends of tip links in frog saccular hair cells. J. Neurosci. 18, 8637–8647.

Gillespie, P. G. and Müller, U., 2009. Mechanotransduction by hair cells: models, molecules, and mechanisms. Cell. 139, 33–44.

Goodyear, R. J., Legan, P. K., Wright, M. B., Marcotti, W., Oganesian, A., Coats, S. A., Booth, C. J., Kros, C. J., Seifert, R. A., Bowen-Pope, D. F. and Richardson, G. P., 2003. A receptor-like inositol lipid phosphatase is required for the maturation of developing cochlear hair bundles. J. Neurosci. 23, 9208–9219.

Hackney, C. M., Fettiplace, R. and Furness, D. N., 1993. The functional morphology of stereociliary bundles on turtle cochlear hair cells. Hear Res. 69, 163–175.

Hasson, T., Gillespie, P. G., Garcia, J. A., MacDonald, R. B., Zhao, Y., Yee, A. G., Mooseker, M. S. and Corey, D. P., 1997. Unconventional myosins in inner-ear sensory epithelia. J. Cell Biol. 137, 1287–1307.

Hirono, M., Denis, C. S., Richardson, G. P. and Gillespie, P. G., 2004. Hair cells require phosphatidylinositol 4,5-bisphosphate for mechanical transduction and adaptation. Neuron. 44, 309–320.

Hutchings, J. and Zanetti, G., 2018. Fine details in complex environments: the power of cryo-electron tomography. Biochem Soc Trans.

Hwang, P., Chou, S. W., Chen, Z. and McDermott, B. M., 2015. The stereociliary paracrystal Is a dynamic cytoskeletal scaffold in vivo. Cell Rep. 13, 1287–1294.

Jacobs, R. A. and Hudspeth, A. J., 1990. Ultrastructural correlates of mechanoelectrical transduction in hair cells of the bullfrog’s internal ear. Cold Spring Harb. Symp. Quant. Biol. 55, 547–561.

Jasnin, M., Asano, S., Gouin, E., Hegerl, R., Plitzko, J. M., Villa, E., Cossart, P. and Baumeister, W., 2013. Three-dimensional architecture of actin filaments in Listeria monocytogenes comet tails. Proc Natl Acad Sci U S A. 110, 20521–20526.

Jontes, J. D. and Milligan, R. A., 1997a. Three-dimensional structure of Brush Border Myosin-I at approximately 20 A resolution by electron microscopy and image analysis. J Mol Biol. 266, 331–342.

Jontes, J. D. and Milligan, R. A., 1997b. Brush border myosin-I structure and ADP-dependent conformational changes revealed by cryoelectron microscopy and image analysis. J. Cell Biol. 139, 683–693.

Kitajiri, S., Fukumoto, K., Hata, M., Sasaki, H., Katsuno, T., Nakagawa, T., Ito, J., Tsukita, S. and Tsukita, S., 2004. Radixin deficiency causes deafness associated with progressive degeneration of cochlear stereocilia. J Cell Biol. 166, 559–570.

Kitajiri, S., Sakamoto, T., Belyantseva, I. A., Goodyear, R. J., Stepanyan, R., Fujiwara, I., Bird, J. E., Riazuddin, S., Riazuddin, S., Ahmed, Z. M., Hinshaw, J. E., Sellers, J., Bartles, J. R., Hammer, J. A., Richardson, G. P., Griffith, A. J., Frolenkov, G. I. and Friedman, T. B., 2010. Actin-bundling protein TRIOBP forms resilient rootlets of hair cell stereocilia essential for hearing. Cell. 141, 786–798.

Kremer, J. R., Mastronarde, D. N. and McIntosh, J. R., 1996. Computer visualization of three-dimensional image data using IMOD. J Struct Biol. 116, 71–76.

Krey, J. F., Krystofiak, E. S., Dumont, R. A., Vijayakumar, S., Choi, D., Rivero, F., Kachar, B., Jones, S. M. and Barr-Gillespie, P. G., 2016. Plastin 1 widens stereocilia by transforming actin filament packing from hexagonal to liquid. J Cell Biol. 215, 467–482.

Liberman, M. C. and Dodds, L. W., 1987. Acute ultrastructural changes in acoustic trauma: serial-section reconstruction of stereocilia and cuticular plates. Hear Res. 26, 45–64.

McIntosh, J. R., O’Toole, E., Morgan, G., Austin, J., Ulyanov, E., Ataullakhanov, F. and Gudimchuk, N., 2018. Microtubules grow by the addition of bent guanosine triphosphate tubulin to the tips of curved protofilaments. J Cell Biol. 217, 2691–2708.

Merritt, R. C., Manor, U., Salles, F. T., Grati, M., Dose, A. C., Unrath, W. C., Quintero, O. A., Yengo, C. M. and Kachar, B., 2012. Myosin IIIB uses an actin-binding motif in its espin-1 cargo to reach the tips of actin protrusions. Curr Biol. 22, 320–325.

Metlagel, Z., Krey, J. F., Song, J., Swift, M. F., Tivol, W. J., Dumont, R. A., Thai, J., Chang, A., Seifikar, H., Volkmann, N., Hanein, D., Barr-Gillespie, P. G. and Auer, M., 2019. Electron cryo-tomography of vestibular hair-cell stereocilia. J Struct Biol. 206, 149–155.

Morgan, C. P., Krey, J. F., Grati, M., Zhao, B., Fallen, S., Kannan-Sundhari, A., Liu, X. Z., Choi, D., Müller, U. and Barr-Gillespie, P. G., 2016. PDZD7-MYO7A complex identified in enriched stereocilia membranes. eLife. 5, e18312. doi: 10.7554/eLife.18312.

Narayanan, P., Chatterton, P., Ikeda, A., Ikeda, S., Corey, D. P., Ervasti, J. M. and Perrin, B. J., 2015. Length regulation of mechanosensitive stereocilia depends on very slow actin dynamics and filament-severing proteins. Nat Commun. 6, 6855.

Oikonomou, C. M. and Jensen, G. J., 2017. Cellular Electron Cryotomography: Toward Structural Biology In Situ. Annu Rev Biochem. 86, 873–896.

Pataky, F., Pironkova, R. and Hudspeth, A. J., 2004. Radixin is a constituent of stereocilia in hair cells. Proc Natl Acad Sci U S A. 101, 2601–2606.

Peng, A. W., Gnanasambandam, R., Sachs, F. and Ricci, A. J., 2016. Adaptation Independent Modulation of Auditory Hair Cell Mechanotransduction Channel Open Probability Implicates a Role for the Lipid Bilayer. J Neurosci. 36, 2945–2956.

Perrin, B. J., Strandjord, D. M., Narayanan, P., Henderson, D. M., Johnson, K. R. and Ervasti, J. M., 2013. β-actin and fascin-2 cooperate to maintain stereocilia length. J Neurosci. 33, 8114–8121.

Pettersen, E. F., Goddard, T. D., Huang, C. C., Couch, G. S., Greenblatt, D. M., Meng, E. C. and Ferrin, T. E., 2004. UCSF Chimera--a visualization system for exploratory research and analysis. J Comput Chem. 25, 1605–1612.

Pontes, B., Monzo, P. and Gauthier, N. C., 2017. Membrane tension: A challenging but universal physical parameter in cell biology. Semin Cell Dev Biol. 71, 30–41.

Powers, R. J., Kulason, S., Atilgan, E., Brownell, W. E., Sun, S. X., Barr-Gillespie, P. G. and Spector, A. A., 2014. The local forces acting on the mechanotransduction channel in hair cell stereocilia. Biophys J. 106, 2519–2528.

Powers, R. J., Roy, S., Atilgan, E., Brownell, W. E., Sun, S. X., Gillespie, P. G. and Spector, A. A., 2012. Stereocilia membrane deformation: implications for the gating spring and mechanotransduction channel. Biophys J. 102, 201–210.

Prost, J., Barbetta, C. and Joanny, J. F., 2007. Dynamical control of the shape and size of stereocilia and microvilli. Biophys J. 93, 1124–1133.

Raucher, D., Stauffer, T., Chen, W., Shen, K., Guo, S., York, J. D., Sheetz, M. P. and Meyer, T., 2000. Phosphatidylinositol 4,5-bisphosphate functions as a second messenger that regulates cytoskeleton-plasma membrane adhesion. Cell. 100, 221–228.

Rehman, A. U., Morell, R. J., Belyantseva, I. A., Khan, S. Y., Boger, E. T., Shahzad, M., Ahmed, Z. M., Riazuddin, S., Khan, S. N., Riazuddin, S. and Friedman, T. B., 2010. Targeted capture and next-generation sequencing identifies C9orf75, encoding taperin, as the mutated gene in nonsyndromic deafness DFNB79. Am J Hum Genet. 86, 378–388.

Roberts, W. M., Howard, J. and Hudspeth, A. J., 1988. Hair cells: transduction, tuning, and transmission in the inner ear. Annu Rev Cell Biol. 4, 63–92.

Robertson, D., Johnstone, B. M. and McGill, T. J., 1980. Effects of loud tones on the inner ear: a combined electrophysiological and ultrastructural study. Hear Res. 2, 39–43.

Rzadzinska, A. K., Schneider, M. E., Davies, C., Riordan, G. P. and Kachar, B., 2004. An actin molecular treadmill and myosins maintain stereocilia functional architecture and self-renewal. J. Cell Biol. 164, 887–897.

Sazzed, S., Song, J., Kovacs, J. A., Wriggers, W., Auer, M. and He, J., 2018. Tracing Actin Filament Bundles in Three-Dimensional Electron Tomography Density Maps of Hair Cell Stereocilia. Molecules. 23,

Schneider, M. E., Dose, A. C., Salles, F. T., Chang, W., Erickson, F. L., Burnside, B. and Kachar, B., 2006. A new compartment at stereocilia tips defined by spatial and temporal patterns of myosin IIIa expression. J Neurosci. 26, 10243–10252.

Sekerkova, G., Zheng, L., Loomis, P. A., Changyaleket, B., Whitlon, D. S., Mugnaini, E. and Bartles, J. R., 2004. Espins are multifunctional actin cytoskeletal regulatory proteins in the microvilli of chemosensory and mechanosensory cells. J Neurosci. 24, 5445–5456.

Sekerkova, G., Zheng, L., Mugnaini, E. and Bartles, J. R., 2006. Differential expression of espin isoforms during epithelial morphogenesis, stereociliogenesis and postnatal maturation in the developing inner ear. Dev Biol. 291, 83–95.

Shin, J. B., Krey, J. F., Hassan, A., Metlagel, Z., Tauscher, A. N., Pagana, J. M., Sherman, N. E., Jeffery, E. D., Spinelli, K. J., Zhao, H., Wilmarth, P. A., Choi, D., David, L. L., Auer, M. and Barr-Gillespie, P. G., 2013. Molecular architecture of the chick vestibular hair bundle. Nat Neurosci. 16, 365–374.

Shin, J. B., Longo-Guess, C. M., Gagnon, L. H., Saylor, K. W., Dumont, R. A., Spinelli, K. J., Pagana, J. M., Wilmarth, P. A., David, L. L., Gillespie, P. G. and Johnson, K. R., 2010. The R109H variant of fascin-2, a developmentally regulated actin crosslinker in hair-cell stereocilia, underlies early-onset hearing loss of DBA/2J mice. J Neurosci. 30, 9683–9694.

Sobin, A. and Flock, A., 1983. Immunohistochemical identification and localization of actin and fimbrin in vestibular hair cells in the normal guinea pig and in a strain of the waltzing guinea pig. Acta Otolaryngol. 96, 407–412.

Steyger, P. S., Gillespie, P. G. and Baird, R. A., 1998. Myosin Iβ is located at tip link anchors in vestibular hair bundles. J. Neurosci. 18, 4603–4615.

Sun, S. Y., Kaelber, J. T., Chen, M., Dong, X., Nematbakhsh, Y., Shi, J., Dougherty, M., Lim, C. T., Schmid, M. F., Chiu, W. and He, C. Y., 2018. Flagellum couples cell shape to motility in Trypanosoma brucei. Proc Natl Acad Sci U S A. 115, E5916–E5925.

Tilney, L. G., DeRosier, D. J. and Mulroy, M. J., 1980. The organization of actin filaments in the stereocilia of cochlear hair cells. J. Cell Biol. 86, 244–259.

Tilney, L. G., Tilney, M. S., Saunders, J. C. and DeRosier, D. J., 1986. Actin filaments, stereocilia, and hair cells of the bird cochlea. III. The development and differentiation of hair cells and stereocilia. Devel. Biol. 116, 100–118.

Tilney, M. S., Tilney, L. G., Stephens, R. E., Merte, C., Drenckhahn, D., Cotanche, D. A. and Bretscher, A., 1989. Preliminary biochemical characterization of the stereocilia and cuticular plate of hair cells of the chick cochlea. J Cell Biol. 109, 1711–23.

Tinevez, J. Y., Perry, N., Schindelin, J., Hoopes, G. M., Reynolds, G. D., Laplantine, E., Bednarek, S. Y., Shorte, S. L. and Eliceiri, K. W., 2017. TrackMate: An open and extensible platform for single-particle tracking. Methods. 115, 80–90.

Turgay, Y., Eibauer, M., Goldman, A. E., Shimi, T., Khayat, M., Ben-Harush, K., Dubrovsky-Gaupp, A., Sapra, K. T., Goldman, R. D. and Medalia, O., 2017. The molecular architecture of lamins in somatic cells. Nature. 543, 261–264.

Vélez-Ortega, A. C., Freeman, M. J., Indzhykulian, A. A., Grossheim, J. M. and Frolenkov, G. I., 2017. Mechanotransduction current is essential for stability of the transducing stereocilia in mammalian auditory hair cells. eLife. 6,

Whittaker, M., Wilson-Kubalek, E. M., Smith, J. E., Faust, L., Milligan, R. A. and Sweeney, H. L., 1995. A 35-A movement of smooth muscle myosin on ADP release [see comments]. Nature. 378, 748–751.

Zhang, D. S., Piazza, V., Perrin, B. J., Rzadzinska, A. K., Poczatek, J. C., Wang, M., Prosser, H. M., Ervasti, J. M., Corey, D. P. and Lechene, C. P., 2012. Multi-isotope imaging mass spectrometry reveals slow protein turnover in hair-cell stereocilia. Nature. 481, 520–524.

Zhao, H., Williams, D. E., Shin, J. B., Brugger, B. and Gillespie, P. G., 2012. Large membrane domains in hair bundles specify spatially constricted radixin activation. J Neurosci. 32, 4600–4609.

Zheng, L., Sekerkova, G., Vranich, K., Tilney, L. G., Mugnaini, E. and Bartles, J. R., 2000. The deaf jerker mouse has a mutation in the gene encoding the espin actin-bundling proteins of hair cell stereocilia and lacks espins. Cell. 102, 377–385.

